# Intranasal vasopressin like oxytocin increases social attention by influencing top-down control, but additionally enhances bottom-up control

**DOI:** 10.1101/2021.05.21.445212

**Authors:** Qian Zhuang, Xiaoxiao Zheng, Benjamin Becker, Wei Lei, Xiaolei Xu, Keith M. Kendrick

**Author notes:** Corresponding authors: Keith M. Kendrick and Xiaolei Xu. Equal contribution (Qian Zhuang, Xiaoxiao Zheng). No. 2006, Xiyuan Ave., West Hi-Tech Zone, Chengdu, Sichuan 611731, China.

## Abstract

The respective roles of the neuropeptides arginine vasopressin (AVP) and oxytocin (OXT) in modulating social cognition and for therapeutic intervention in autism spectrum disorder have not been fully established. In particular, while numerous studies have demonstrated effects of oxytocin in promoting social attention the role of AVP has not been examined. The present study employed a randomized, double-blind, placebo (PLC)-controlled between-subject design to explore the social- and emotion-specific effects of AVP on both bottom-up and topdown attention processing with a validated emotional anti-saccade eye-tracking paradigm in 80 male subjects (PLC = 40, AVP = 40). Our findings showed that AVP increased the error rate for social (angry, fearful, happy, neutral and sad faces) but not non-social (oval shapes) stimuli during the anti-saccade condition and reduced error rates in the pro-saccade condition. Comparison of these findings with a previous study (sample size: PLC = 33, OXT = 33) using intranasal oxytocin revealed similar effects of the two peptides on anti-saccade errors but a significantly greater effect of AVP on pro-saccades. Both peptides also produced a post-task anxiolytic effect by reducing state anxiety. Together these findings suggested that both AVP and OXT decrease goal-directed top-down attention control to social salient stimuli but that AVP more potently increased bottom-up social attentional processing.

## 1. Introduction

A large body of research in both animal models and humans has demonstrated important modulatory effects of the hypothalamic neuropeptides arginine vasopressin (AVP) and oxytocin (OXT) on social cognition (Abramova et al., 2020; Albers, 2012; Caldwell, 2017). In contrast to their distinct peripheral physiological effects following release from the posterior pituitary these two highly similar and evolutionary conserved peptides tend to have broadly similar effects on social cognition and bonds when acting on their receptors in the brain (Abramova et al., 2020; Albers, 2012; Caldwell, 2017; Gozzi et al., 2017) and clinical trials involving chronic treatment with intranasal AVP and OXT in autistic children have also produced equivalent improvements in social responsivity (Parker et al., 2017, 2019). Nevertheless, there is some evidence that they can produce different effects on brain circuits (Galbusera et al., 2017) and notably that they may have even opposing effects on amygdala responses (Kou et al., 2020; Lieberz et al., 2020) and on anxiety and depression (Neumann and Landgraf, 2012). There may also be differences between the functional effects of the two peptides in the context of social decision making, cooperation and empathy (Neto et al., 2020; Rilling et al., 2014; Tabak et al., 2015; Yang et al., 2021) and AVP has an established role in promoting male aggression in animal models (Ferris, 2005) and in terms of angry facial expressions in human males (Thompson et al., 2004, 2006).

One of the key fundamental effects of OXT in the context of social cognition is its facilitation of attention towards social cues (Le et al., 2020; Xu et al., 2015; Xu et al., 2019; Yao et al., 2018) and this may in turn contribute to its modulatory effects on many different aspects of social recognition memory and other key social behaviors. While some studies in humans have also reported similar effects of AVP on social recognition memory, social cooperation and decision making (Abramova et al., 2020; Albers, 2012; Caldwell, 2017; Guastella et al., 2010, 2011; Zink et al., 2011), the specific role of this peptide in modulating social attention in humans has not been established, although an early study in rats has reported effects of an AVP metabolite on selective attention (Bunsey et al., 1990).

The anti-saccade task has been used extensively to examine both bottom-up and topdown attention control (Hallett, 1978; Munoz and Everling, 2004). In this task participants are either instructed to shift their attention towards a stimulus by a saccadic eye movement (prosaccade) or to make a saccade in the opposite direction (anti-saccade). Pro-saccades require the engagement of bottom-up attention control to focus on the target automatically, while antisaccades require top-down attention control to prevent reflexive orienting towards salient stimuli (Munoz and Everling, 2004). The engagement of the effortful cognitive control demand is also reflected in longer latencies and more errors during anti-saccade as compared to pro-saccade conditions (Hutton and Ettinger, 2006; Munoz and Everling, 2004). In a previous study we used a validated emotional anti-saccade task with social (happy, angry, fearful, sad, and neutral faces) and non-social stimuli (oval shapes) to demonstrate that intranasal OXT selectively increases anti-saccade errors towards social stimuli in this task indicating that it may increase implicit attention towards them by reducing the effects of topdown attention control (Xu et al., 2019). Additionally, we found that OXT reduced post-task state anxiety scores in support of its anxiolytic effects also found in other contexts (Kou et al., 2020; Neumann and Landgraf, 2012). We could therefore use this same task to determine whether AVP produces similar effects on top-down attention control as OXT and/or also on bottom-up control.

Against this background, the present study combined the anti-saccade eye-tracking paradigm we used in a previous study to establish modulatory effects of intranasal OXT (Xu et al., 2019) to explore common and distinguishable effects of intranasal AVP on bottom-up attention processing (pro-saccade errors) or top-down attention control (anti-saccade errors) as well as anxiety in male subjects. The present findings were then compared with our previous randomized pre-registered study in which male subjects underwent intranasal OXT administration (NCT03486925 – Xu et al., 2019) using the same spray device, treatment timing protocol and task paradigm. Given the background of previous reported similarities in effects of OXT and AVP on aspects of social recognition memory (Abromova et al., 2020; Zink et al., 2011), social responsivity in autism (Parker et al., 2017, 2019) and opposing effects on anxiety (Neumann and Landgraf, 2012), we hypothesized that AVP would produce similar increased attention allocation towards social salient stimuli as OXT, resulting in either, or both, increased anti-saccade and decreased pro-saccade errors. On the other hand, we hypothesized that AVP would increase rather than decrease associated state anxiety.

## 2. Material and methods

### 2.1. Participants

80 healthy male subjects (age: Mean ± SEM = 21.93 ± 0.25 years) were recruited from the University of Electronic Science and Technology of China (UESTC). Only male subjects were recruited for the study in order to match our previous one investigating effects of intranasal OXT (Xu et al., 2019). All subjects were instructed to abstain from caffeine and alcohol 24 hours before the experiment. None of them self-reported previous or current psychiatric or neurological disorders. Exclusion criteria were regular medication, illicit or licit substance use including nicotine. In the double-blind, placebo-controlled between-subject design experiment, all subjects were randomly assigned to receive 20 International Units (IU) of intranasal AVP (n = 40, age: Mean ± SEM = 21.18 ± 0.30 years) or placebo (PLC) spray (n = 40, age: Mean ± SEM = 22.68 ± 0.38 years). Intranasal AVP (AVP supplied by Bio-Techne China Co., Ltd) was administered dissolved in identical ingredients (sodium chloride and glycerol) as in previous experiments using OXT nasal spray and administered sterile (0.22μm Millipore filtered - six 0.1ml puffs, 3 to each nostril and interspaced by 30 s) 45 min prior to the start of the anti-saccade paradigm. The sterile PLC spray was identical in composition other than the AVP (supplied by Sichuan Meike, Pharmaceutical Company, Sichuan, China). A 20IU AVP dose was chosen since it is equivalent to 24IU of OXT used in our previous study (Xu et al., 2019) and the same as that used in a number of previous studies (Alyousefi-van Dijk et al., 2019; Neto et al., 2020; Pietrowsky et al., 1996; Tabak et al., 2015). See **supplementary Figure S1** for consolidated standards of reporting trials (CONSORT) flow chart.

The present study was approved by the local ethics committee of University of Electronic Science and Technology of China (UESTC) and in accordance with the latest version of the Declaration of Helsinki and pre-registered at clinical trials.gov (ClinicalTrials.gov ID: NCT04493554). All subjects signed written informed consent and received monetary compensation after the experiment (130 RMB).

### 2.2. Procedures

To control for the potential confounding effects of between-subject differences in mood and personality traits, subjects were required to complete validated psychometric questionnaires in Chinese before the AVP or PLC treatment including Positive and Negative Affect Schedule (PANAS, Watson et al., 1988), State-Trait Anxiety Inventory (STAI, Spielberger, 1983) and some other scales (details see **Supplementary Material)**. Next, subjects were randomized to receive either 20 IU AVP or PLC intranasal spray 45 min before the eye-tracking task. Additionally, subjects were asked to complete PANAS and STAI questionnaires before and after the experimental session to further examine the potential effects of AVP treatment and the task on mood and state anxiety. After the whole experimental task, subjects could not accurately identify which treatment (AVP/PLC) they had received with their guesses not being significant different from chance (*χ*^2^ = 2.69, *p* = 0.10).

### 2.3. Experimental Task

The present study used a validated anti-saccade paradigm (Chen et al., 2014; Xu et al., 2019) including five social-emotional faces (angry, sad, happy, fearful and neutral) from 8 actors (4 male and 4 female actors) and 8 non-social slightly varying oval shapes. To avoid any carryover effect of emotional stimuli, the task always started with 2 non-social blocks (one anti- and one pro-saccade block) including 48 trials in each block followed by 12 emotional blocks (6 anti- and 6 pro-saccade blocks) including 40 trials per block and 8 trials per emotion. The anti- and pro-saccade blocks and trials within each block were presented randomly. Each block started with a 2000 ms cue word “Towards” or “Away” as instructions (**Figure 1**). Following the cue, a jittered fixation appeared with a mean time duration of 1500 ms (time range:1000 ms-3500 ms) and subjects were asked to fixate on it. Next, the stimulus appeared at the left or right side of the screen for 1000 ms. Subjects were asked to look at the opposite side to the stimulus during the “Away” blocks (anti-saccade condition) and look towards the stimulus during the “Towards” blocks (pro-saccade condition) as quickly and accurately as possible. The task took 35 min to complete with a brief rest for subjects between blocks.

**Figure 1.**
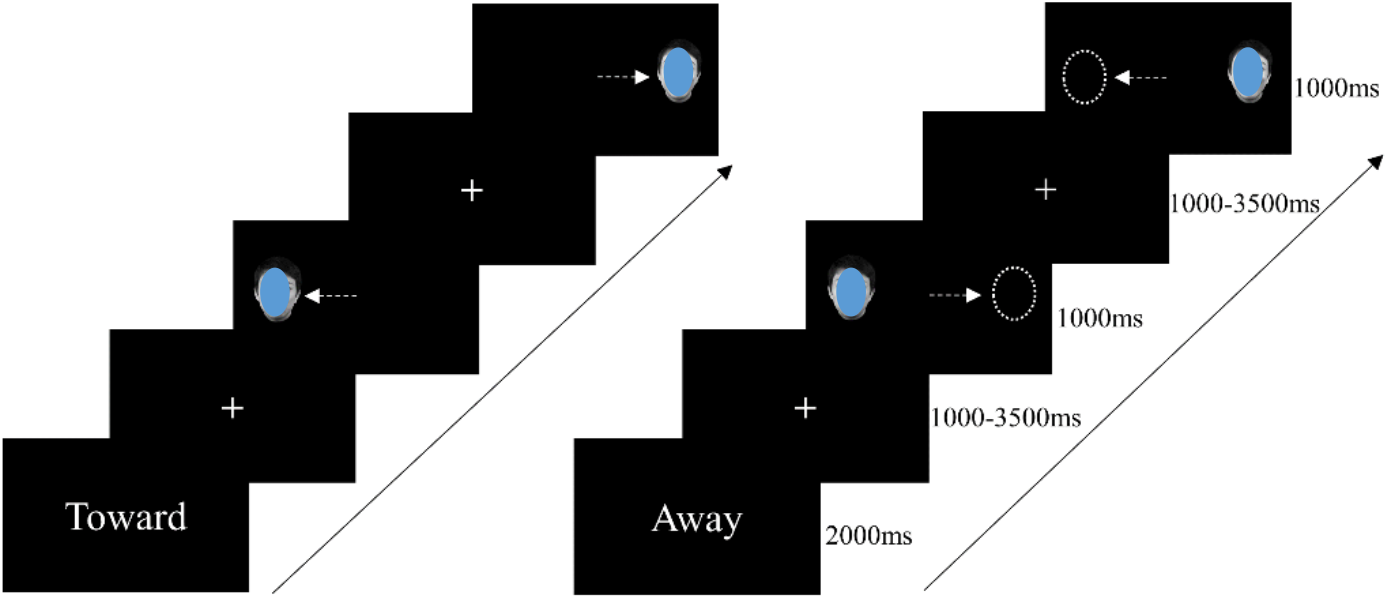
The emotional anti-saccade paradigm and timing set including the prosaccade condition (left) and anti-saccade condition (right). Note: for the preprint the face has been overlayed with an oval shape.

### 2.4. Eye tracking data Recording and Processing

Eye movement data were recorded using an EyeLink 1000 Plus system (SR Research, Ottawa, Canada) sampling at 2000 Hz with a screen resolution of 1024*768. A chin rest 57cm away from the monitor was used to ensure standard distance and position from the screen. Before each block a 9-point calibration was performed to assure eye-tracking data quality.

The raw eye-tracking data were preprocessed and exported by the EyeLink DataViewer 3.1 (SR Research Mississauga, Ontario, Canada). In accordance with previous studies, trials were discarded with latencies less than 70 ms or more than 700 ms, and saccade velocity lower than 30°/sec (Xu et al., 2019). Subjects with more than 25% of their total trial number deleted were also excluded (Reinholdt-Dunne et al., 2012). Finally, only 2 subjects were excluded from the final analyses due to technical error. There was no significant difference in percentage of excluded trials for the remaining subjects between AVP (n = 39) and PLC groups (n = 39, AVP: Mean ± SEM = 8.14% ± 0.64%, PLC: Mean ± SEM = 9.18% ± 0.88%, t = 0.96, *p* = 0.34). The mean error rate and correct saccade latency during both anti- and prosaccade conditions were calculated and served as dependent variables.

### 2.5. Statistical analyses

To explore the social-specific effects of AVP on attention control, 2 (treatment: AVP/PLC)*2 (condition: social/non-social)*2 (task: pro-/anti-saccade) mixed ANOVAs were performed on both latency and error rate. To further investigate the emotion-specific effects of AVP on attention processing, we additionally conducted 2 (treatment: AVP/PLC)*2 (task: pro-/anti-saccade)*6 (stimuli: angry/sad/fearful/happy/neutral/shape) repeated-measures ANOVAs on latency and error rate.

To examine the differences between effects of AVP and OXT on attention processing, 3 (treatment: AVP/OXT/PLC)*2 (task: pro-/anti-saccade)*2 (condition: social/non-social) mixed ANOVAs were conducted on latency and error rate. Additionally, to further control for non-treatment related factors the two intranasal PLC groups from the current (PLC) and previous study (PLC2, ClinicalTrials.gov ID: NCT03486925) were compared with respect to the social- and emotion-specific effects on attentional control. The comparison study was preregistered at clinical trials.gov (ClinicalTrials.gov ID: NCT04715737). Details see **Supplementary Material**. Appropriate Bonferroni correction for multiple comparisons was employed to disentangle significant main and interaction effects.

## 3. Results

### 3.1. Potential confounders

Independent t-tests were used to examine the potential confounding effect of mood and personality for questionnaire scores. Results showed no significant differences in mood and personality traits between the PLC and AVP group before and after treatment (all *ps* > 0.17, **details see Supplementary Material Table S1**). Paired t-tests were conducted to examine treatment effects on post-task anxiety measured by SAI and revealed that scores were significantly decreased during post-task in the AVP group (pre: Mean ± SEM = 40.05 ± 1.43, post: Mean ± SEM = 37.62 ± 1.09, *p* = 0.004, Cohen’s *d* = 0.31) but not in the PLC group (pre: Mean ± SEM = 38.77 ± 1.47, post: Mean ± SEM = 38.28 ± 1.47, *p* = 0.59). This suggested that the intranasal AVP treatment had an anxiolytic effect.

### 3.2. Response latency

The 2 (treatment: AVP/PLC)*2 (condition: social/non-social)*2 (task: pro-/anti-saccade) mixed ANOVA on latency showed a significant main effect of task (F_1, 76_ = 872.11, *p* < 0.001, η_p_^2^ = 0.92) with longer anti-saccade than pro-saccade latencies (anti: Mean ± SEM = 263.87 ± 2.91 ms, pro: Mean ± SEM = 190.56 ± 2.46 ms) and a main effect of condition (F_1, 76_ = 119.94, *p* < 0.001, η_p_^2^ = 0.61) with faster latencies for social than non-social stimuli (social: Mean ± SEM = 219.94 ± 2.62 ms, non-social: Mean ± SEM = 234.48 ± 2.34 ms).

Additionally, a significant interaction between task and condition was observed (F_1, 76_ = 90.25, *p* < 0.001, η_p_^2^ = 0.54). Post-hoc Bonferroni-corrected comparisons indicated that latencies for social stimuli were significantly faster than non-social stimuli during pro-saccade but not antisaccade condition (pro-saccade: social: Mean ± SEM = 175.91 ± 2.42 ms, non-social: Mean ± SEM = 205.21 ± 2.77 ms, *p* < 0.001; anti-saccade: social: Mean ± SEM = 263.98 ± 3.31 ms, non-social: Mean ± SEM = 263.76 ± 2.96 ms, *p* = 0.93). No other significant interaction effects between treatment, condition and task were found (all *ps* > 0.06).

To further explore emotion-specific effects of AVP on attentional processing, a 2 (treatment: AVP/PLC)*2 (task: pro-/anti-saccade)*6 (stimuli: angry/sad/fearful/happy/ neutral/shape) repeated-measures ANOVA was conducted. Results showed a main effect of task (F_1, 76_ = 1164.52, *p* < 0.001, η_p_^2^ = 0.94) reflecting faster latencies in the pro-saccade than anti-saccade condition (anti: Mean ± SEM = 262.87 ± 2.96 ms, pro: Mean ± SEM = 180.00 ± 2.08 ms) and a main effect of stimuli (F_5, 380_ = 79.84, *p* < 0.001, η_p_^2^ = 0.51) reflecting longer latencies for shapes than all faces (all *ps* < 0.001). Moreover, we observed a significant interaction between task and stimuli (F_5, 380_ = 52.58, *p* < 0.001, η_p_^2^ = 0.41). Post-hoc Bonferroni-corrected comparisons showed that the response latencies for shapes were longer than those for all faces during the pro-saccade (all *ps* < 0.001) but not anti-saccade condition (all *ps* > 0.99). Finally, a three-way (task * stimuli * treatment) interaction effect was significant (F_5, 380_ = 3.42, *p* = 0.02, η_p_^2^ = 0.04) and post-hoc Bonferroni-corrected comparisons indicated that the interaction was driven by the main effects of task and stimuli (task: all *ps* < 0.001, longer latencies for anti-compared to pro-saccade; stimuli: all *ps* < 0.001, longer latencies for shapes compared to all faces in pro-saccade condition; treatment: all *ps* > 0.07).

### 3.3. Error rate

A mixed ANOVA on error rate revealed a significant main effect of task (F_1, 76_ =130.09, *p* < 0.001, η_p_^2^ = 0.63) with higher error rates in the anti-saccade than pro-saccade condition (anti: Mean ± SEM = 10.92% ± 0.83%, pro: Mean ± SEM = 1.65% ± 0.25%) and a main effect of condition (F_1, 76_ =28.82, *p* < 0.001, η_p_^2^ = 0.28) with higher error rates for social compared to non-social stimuli (social: Mean ± SEM = 7.57% ± 0.53%, non-social: Mean ± SEM = 5.00% ± 0.51%). Significant interactions between condition and task (F_1, 76_ =29.66, *p* < 0.001, η_p_^2^ = 0.28) and task and treatment (F_1, 76_ =6.12, *p* = 0.016, η_p_^2^ = 0.08) were also observed. Post-hoc Bonferroni-corrected comparisons indicated increased error rates for social compared to nonsocial stimuli during the anti-saccade (social: Mean ± SEM = 13.62% ± 0.98%, non-social: Mean ± SEM = 8.21% ± 0.91%, *p* < 0.001, Cohen’s *d* = 0.63) but not pro-saccade condition (social: Mean ± SEM = 1.53% ± 0.28%, non-social: Mean ± SEM = 1.78% ± 0.37%, *p* = 0.53). Compared to PLC, AVP increased error rates on anti-saccade condition (PLC: Mean ± SEM = 9.18% ± 1.17%, AVP: Mean ± SEM = 12.66% ± 1.17%, *p* = 0.039, Cohen’s *d* = 0.48) but not pro-saccade condition (PLC: Mean ± SEM = 1.92% ± 0.36%, AVP: Mean ± SEM = 1.38% ± 0.36%, *p* = 0.29). Moreover, the treatment * task * condition interaction was significant (F_1, 76_ =7.48, *p* = 0.008, η_p_^2^ = 0.09) and post-hoc Bonferroni-corrected comparisons revealed that compared to PLC treatment, AVP significantly increased error rates for social but not non-social stimuli in anti-saccade condition (social: PLC: Mean ± SEM = 10.79% ± 1.39%, AVP: Mean ± SEM = 16.46% ± 1.39%, *p* = 0.005, Cohen’s *d* = 0.36; non-social: PLC: Mean ± SEM = 7.57% ± 1.29%, AVP: Mean ± SEM = 8.85% ± 1.29%, *p* = 0.48, **Figure 2A**). In the pro-saccade condition AVP decreased error rates for social rather than non-social stimuli (social: PLC: Mean ± SEM = 2.12% ± 0.39%, AVP: Mean ± SEM = 0.93% ± 0.39%, *p* = 0.034, Cohen’s *d* = 1.40; non-social: PLC: Mean ± SEM = 1.72% ± 0.52%, AVP: Mean ± SEM = 1.83% ± 0.52%, *p* = 0.88).

**Figure 2.**
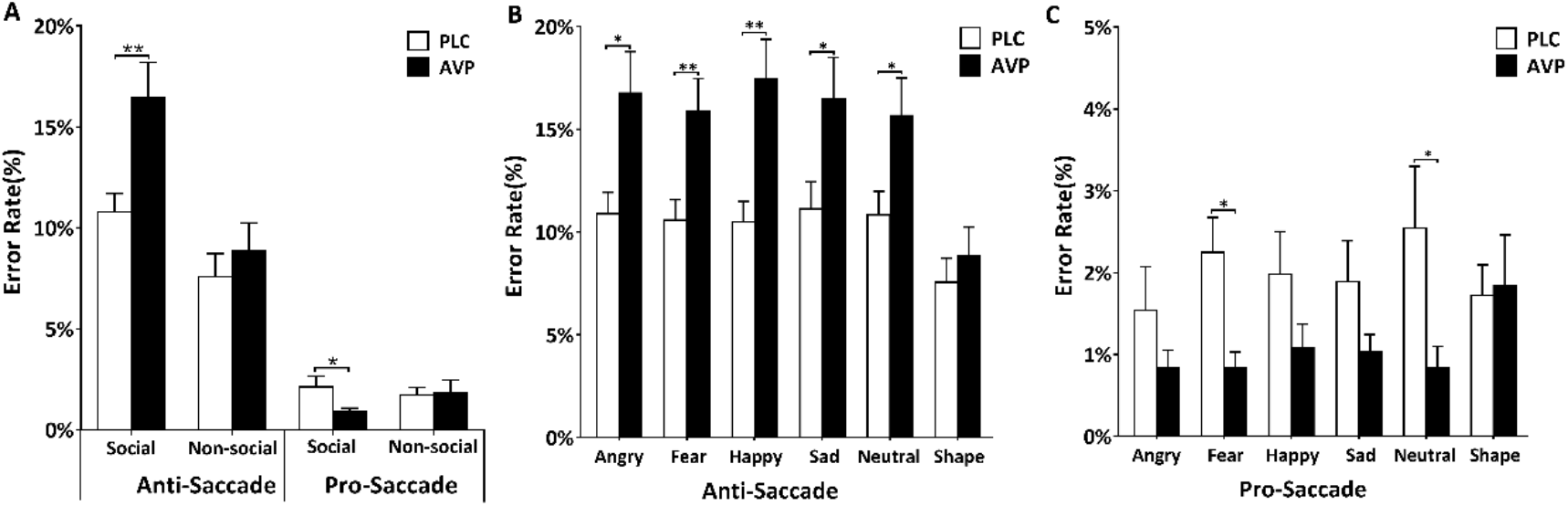
Intranasal AVP’s effect on error rates. (A) Error rates for social and nonsocial stimuli during pro- and anti-saccade condition after AVP and PLC treatment. (B) Error rates for emotional faces and oval shapes during the anti-saccade condition after AVP and PLC treatment (C) Error rates for emotional faces and oval shapes during the pro-saccade condition after AVP and PLC treatment. * and ** denote significant post-hoc difference at *p_Bonferroni_* < 0.05 and *p_Bonferroni_* < 0.01 respectively.

Furthermore, a 2 (treatment: AVP/PLC)*2 (task: pro-/anti-saccade)*6 (stimuli: angry/ sad/fearful/happy/neutral/shape) repeated-measures ANOVA was conducted to explore emotion-specific effects of AVP on error rate. Results revealed a significant main effect of stimuli (F_5, 380_ =11.29, *p* < 0.001, η_p_^2^ = 0.13) with higher error rates for faces compared to shapes (all*ps* < 0.001) and a main effect of task (F_1, 76_ =156.67, *p* < 0.001, η_p_^2^ = 0.67) with increased error rates for the anti-saccade compared to pro-saccade condition (anti: Mean ± SEM = 12.72% ± 0.91%, pro: Mean ± SEM = 1.53% ± 0.23%). Additionally, the interactions between stimuli and task (F_5, 380_ =12.15, *p* < 0.001, η_p_^2^ = 0.14) as well as treatment and task (F_1, 76_ =10.70, *p* = 0.002, η_p_^2^ = 0.12) were significant. Post-hoc analysis with Bonferroni correction showed that compared to the PLC group, AVP increased error rates during the antisaccade condition (PLC: Mean ± SEM = 10.25% ± 1.28%, AVP: Mean ± SEM = 15.19% ± 1.28%, *p* = 0.008, Cohen’s *d* = 0.62) and decreased them during pro-saccade condition (PLC: Mean ± SEM = 1.99% ± 0.33%, AVP: Mean ± SEM = 1.08% ± 0.33%, *p* = 0.05, Cohen’s *d* = 0.72). Error rates for faces were significantly higher compared to those for shapes in the antisaccade (all *ps* < 0.001) but not pro-saccade condition (all *ps* > 0.999). Interestingly, results showed a significant three-way interaction effect between treatment, task and stimuli (F_5, 380_ = 3.01, *p* = 0.021, η_p_^2^ = 0.04). Post-hoc Bonferroni-corrected comparisons revealed higher error rates for all face emotions (faces: all *ps* < 0.029, effect sizes: angry: Cohen’s *d* = 0.58, sad: Cohen’s *d* = 0.51, fearful: Cohen’s *d* = 0.63, happy: Cohen’s *d* = 0.73, neutral: Cohen’s *d* = 0.51) than shapes *(p* = 0.48, **Figure 2B**) after AVP compared to PLC treatment in the antisaccade condition but lower error rates only for fearful and neutral faces (all *ps* < 0.036, effect sizes: fearful: Cohen’s *d* = 0.68, neutral: Cohen’s *d* = 0.48, **Figure 2C**) in the pro-saccade condition.

### 3.4. The differences between effects of intranasal OXT and AVP on attention control

To compare the effects of OXT and AVP on attention control, a 3 (treatment: AVP/OXT/ PLC)*2 (task: pro-/anti-saccade)*2 (condition: social/non-social) repeated-measures ANOVA on latency was performed. However, no significant differences between the effects of OXT and AVP were found (all *ps* > 0.22).

For error rate, a mixed ANOVA analysis revealed a significant three-way interaction between treatment, condition and task (F_2, 108_ =5.12, *p* = 0.008, η_p_^2^ = 0.09). The direct comparison between OXT and AVP’s effects with post-hoc Bonferroni-corrected comparisons showed higher error rates for social stimuli in the pro-saccade condition following OXT compared to AVP treatment (OXT: Mean ± SEM = 3.61% ± 0.51%, AVP: Mean ± SEM = 0.93% ± 0.47%, *p* < 0.001, Cohen’s *d* = 0.96, **Figure 3**). This indicated that AVP enhanced subjects’ automatic bottom-up attention towards social stimuli compared to OXT treatment. There was no significant difference between OXT and AVP for anti-saccade error rates (social: OXT: Mean ± SEM = 21.43% ± 1.72%, AVP: Mean ± SEM = 16.46% ± 1.59%, *p* = 0.11; non-social: OXT: Mean ± SEM = 12.06% ± 1.46%, AVP: Mean ± SEM = 8.86% ± 1.34%, *p* = 0.33), indicating that they had similar effects on top-down attention control.

**Figure 3.**
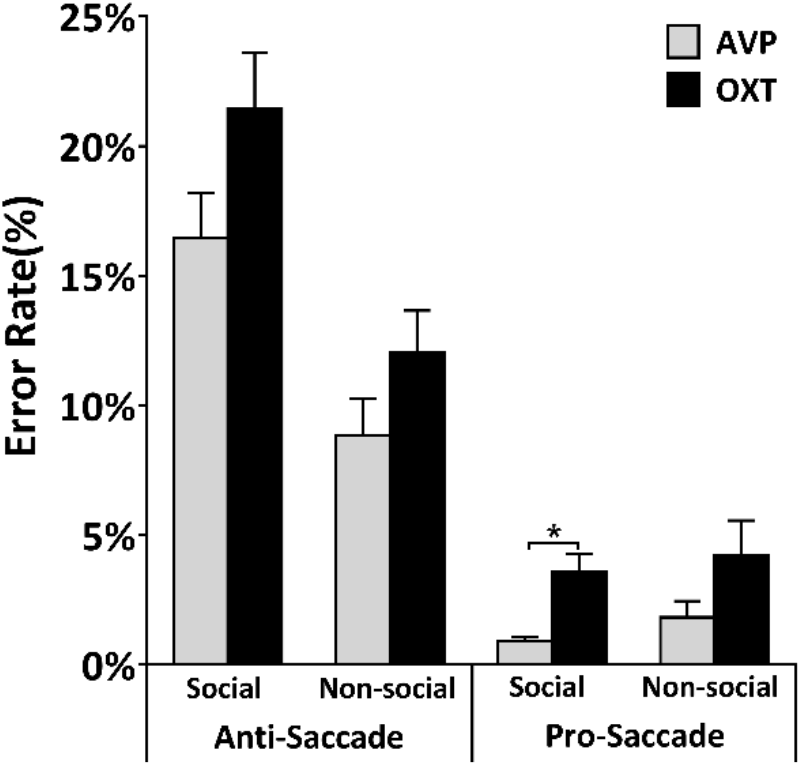
The comparison between AVP and OXT’s social effect on error rates. Post-hoc Bonferroni-corrected comparisons showed higher error rates for social stimuli during pro-saccade condition after OXT treatment compared to AVP treatment. * denotes significant post-hoc difference at *p_Bonferroni_* < 0.05.

### 3.5. Comparison between intranasal OXT and AVP’s effect on state anxiety

To compare intranasal OXT and AVP’s effect on state anxiety, a 3 (group: AVP/OXT/ PLC)*2 (SAI: pre-/post-test) mixed ANOVA was conducted on SAI scores. However, neither the interaction effect (F_2, 108_ = 1.82, *p* = 0.17) nor the main effect of group was significant (F_2, 108_ = 1.50, *p* = 0.23). This suggests that both OXT and AVP attenuated anxiety to a comparable extent.

## 4. Discussion

The current pharmacological eye tracking study used an emotional anti-saccade task to examine the effects of intranasal AVP on both goal-directed top-down attention control and stimulus-driven bottom-up attention processing. Our findings showed that AVP increased error rates for social stimuli during the anti-saccade condition but decreased them during the pro-saccade one. In an additional comparison between the current observed effects of AVP with those for OXT in a previous study (Xu et al., 2019) the two peptides showed equivalent potency in their influence on anti-saccade errors towards face expressions but AVP was significantly more potent than OXT in decreasing pro-saccade ones. Both peptides also had equivalent effects in reducing post-task state anxiety scores. Together these findings indicate that AVP decreases goal-directed top-down attention control to social salient stimuli and increases that of bottom-up control. In contrast to OXT, AVP produced a greater effect on bottom-up attention during processing of social stimuli but the two peptides had equivalent effects on reducing top-down attention and anxiety.

The results showing faster response latencies and higher error rates for social compared to non-social stimuli and longer response latencies and increased error rates in anti-saccade compared to pro-saccade condition are consistent with previous findings that social stimuli decrease the performance especially in the anti-saccade condition and further validate this attentional processing paradigm (Salvia et al., 2020; Xu et al., 2019).

Our findings that AVP increased error rates to social stimuli (face expressions) but not shapes during the anti-saccade condition and also reduced error rates during the pro-saccade condition indicates that it both reduced subjects’ ability to disengage attention away from social stimuli and also increased their attention to them when required to do so. This provides the first clear evidence that AVP can enhance attention to social stimuli in humans and suggests that it can do so both via modulation of top-down and bottom-up processing. It also provides support for a previous study in rats reporting that a fragment of AVP enhanced selective attention to reward cues (Bunsey et al., 1990). Thus previous reported effects of AVP in facilitating social recognition memory and other aspects of social cognition (Abromova et al., 2020; Albers, 2012; Caldwell, 2017; Guastella et al., 2010) may be at least in part contributed to by its modulation of social attention processing.

The influence of AVP in increasing anti-saccade errors occurred across all face expressions although the emotion-specific analysis for pro-saccade errors only revealed significant effects for fearful and neutral expression faces. In general this would indicate that both top-down and bottom-up effects of AVP on social attention processing are to some extent valence independent. Previous studies reporting effects of AVP on recognition memory or responses to emotional faces have been somewhat inconsistent with both valence independent findings (Guastella et al., 2010) as well ones reporting an influence on either negative (Thompson et al., 2004, 2006; Uzefovsky et al., 2012) or positive emotions (Wu et al., 2018). One study using the reading the mind in the eyes test has also failed to find any influence of AVP on recognition of either positive or negative emotions from the eye region (Kenyon et al., 2013). Possibly therefore while AVP may generally facilitate attention towards social stimuli such as faces, its impact on memory and responses to social stimuli may be context dependent.

A comparison of the respective effects of AVP and OXT on this same attentional processing task revealed that the two peptides have broadly similar valence independent effects on increasing attention towards social but not non-social stimuli. Broadly both peptides seem to achieve this social attention bias by disengaging top-down attention to other competing stimuli. In the case of AVP this may be in part due to increased bottom-up attention on social stimuli acting to interfere with competing top-down control. Interestingly, AVP appears to have a stronger influence on bottom-up attentional processing than OXT which possibly indicates that it has more of a dual role in both reducing attention towards competing non-social stimuli as well as increasing automatic attention towards social stimuli. It could be thus argued in the same way as it has been for OXT (Shamay-Tsoory and Abu-Akel, 2016), that AVP is generally promoting the salience of social stimuli. Under the circumstances it is understandable that the two peptides have a number of similar effects on social cognition (Abromova et al., 2020; Albers, 2012; Caldwell, 2017; Gozzi et al., 2017) and importantly in terms of improving social responsivity dysfunction in children with autism (Parker et al., 2017; 2019). Indeed, marked impairments in attentional processing of social stimuli have been reported in individuals with autism spectrum disorders (Amso et al., 2014; Dawson et al., 1998; Kou et al., 2019), particularly in the context of competing demands in top-down attention (Wang et al., 2014). Interestingly too, despite some evidence for opposite effects of OXT and AVP on anxiety (Neumann and Landgraf., 2012) at least in the context of the current paradigm we found that both had a slight but significant anxiolytic effect in reducing posttask state anxiety scores.

In terms of the neural mechanisms whereby AVP and OXT may exert common effects on attention the most likely are the amygdala and insula and their connectivity. For example, several studies have reported that both AVP (Brunnlieb et al., 2013) and OXT (Gao et al., 2016; Luo et al., 2017) increase functional connectivity between the insula or anterior cingulate and amygdala during face processing tasks. In the case of OXT a study on resting state effective connectivity has demonstrated that it primarily strengthens top-down connectivity between frontal, salience, social cognition and reward networks with the amygdala (Jiang et al., 2021). Possibly, AVP may also influence bottom-up connectivity from the amygdala to salience and frontal networks although this has yet to be investigated.

There are several limitations in the current study. First, although the social stimuli we used included male and female faces, we did not explore AVP’s effect on attention control to same- and other-sex faces. Given that previous findings have shown the effects of intranasal AVP are modulated by the sex of emotional faces (Rilling et al., 2017; Wu et al., 2019), it is worth examining the sex-specific effect of AVP on attention processing in the future studies. Second, oval shapes served as non-social stimuli in the present study and these stimuli had a lower complexity compared to emotional faces. Since AVP’s has been reported to have modulatory effects on visual information processing (Wacker and Ludwig, 2019), future studies could consider using natural scenes or objects as non-social control stimuli to match the complexity of faces. Finally, the current study only included male subjects in line with the previous OXT one (Xu et al., 2019). There is increasing evidence for sex-dependent effects of both AVP and OXT (Gao et al., 2016; Luo et al., 2017; Rilling et al., 2017) and so it is possible that the peptides may have different effects on attentional processing in females.

In conclusion, the current study using an emotional anti-saccade task demonstrated that intranasal AVP increases attention towards social stimuli by decreasing effects of competing goal-directed top-down attention and also by increasing the influence of bottom-up processing. In comparison with OXT, AVP had similar effects on top-down attention processing and for reducing anxiety but was more potent than OXT in influencing bottom-up mediated attentional effects.

## Declaration of conflicting interests

The authors declared no conflicts of interest with their research, authorship or the publication of this article.

## Funding

This work was supported by National Natural Science Foundation of China (NSFC) (grant numbers 31530032 – KMK and 91632117 - BB), Key Technological Projects of Guangdong Province (grant number 2018B030335001 – KMK) and Applied & Basic Research Project of Southwest Medical University (grant number 2017-ZRZD-006).

## Notes

### Competing Interest Statement

The authors have declared no competing interest.

