## Supplemental Material for "Intranasal vasopressin like oxytocin increases social attention by influencing top-down control, but additionally enhances bottom-up control"

**Comparison between two PLC groups**

To further control for non-treatment related factors the two intranasal PLC groups (PLC/PLC2, PLC from current AVP study and PLC2 from previous OXT study) was compared with respect to the social- and emotion-specific effects on attentional control.

**Comparison between two PLC groups on latency**

A 2 (group: PLC/PLC2)*2 (condition: social/non-social)*2 (task: pro-/anti-saccade) mixed ANOVA on response latency was conducted to examine the social effects on attention processing between two PLC groups. Results showed a significant main effect of task ($F_{1, 70}$ = 783.68, $p$ < 0.001, $\eta_p^2$ = 0.92), with increased latencies for anti-saccade compared to pro-saccade condition (anti: Mean ± SEM = 271.75 ± 3.07 ms, pro: Mean ± SEM = 190.71 ± 2.30 ms) and a main effect of condition ($F_{1, 70}$ = 79.78, $p$ < 0.001, $\eta_p^2$ = 0.53) with faster latencies for social than non-social stimuli (social: Mean ± SEM = 223.71 ± 2.45 ms, non-social: Mean ± SEM = 238.76 ± 2.43 ms). Additionally, a significant interaction between task and condition was observed ($F_{1, 70}$ = 42.21, $p$ < 0.001, $\eta_p^2$ = 0.38). Post-hoc Bonferroni-corrected comparisons indicated significantly faster latencies for social than non-social stimuli during pro-saccade condition (social: Mean ± SEM = 177.98 ± 2.35 ms, non-social: Mean ± SEM = 203.45 ± 2.54

ms, $p < 0.001$) but not during anti-saccade condition (anti-saccade: social: Mean ± SEM = 269.44 ± 3.28 ms, non-social: Mean ± SEM = 274.06 ± 3.48 ms, $p = 0.11$).

To further explore whether there were significant differences between two PLC groups in the emotion-specific effects on attention control, a 2 (group: PLC/PLC2)*2 (task: pro-/anti-saccade)*6 (stimuli: angry/sad/fearful/happy/neutral/shape) repeated-measures ANOVA on latency was conducted. Results showed a main effect of stimuli ($F_{5, 350} = 68.90$, $p < 0.001$, $\eta_p^2 = 0.50$) and a main effect of task ($F_{1, 70} = 946.18$, $p < 0.001$, $\eta_p^2 = 0.93$), with faster pro-saccade latencies than anti-saccade latencies (anti: Mean ± SEM = 268.86 ± 2.96 ms, pro: Mean ± SEM = 181.05 ± 1.87 ms) and longer latencies for shapes compared to emotional faces (all *ps* < 0.001). Additionally, a significant interaction between task and stimuli was observed ($F_{5, 350} = 23.74$, $p < 0.001$, $\eta_p^2 = 0.25$). Post-hoc Bonferroni-corrected tests showed that the latency for shapes was significantly longer than faces during pro-saccade condition (all *ps* < 0.001) but not anti-saccade condition (all *ps* > 0.086).

**Comparison between two PLC groups on error rate**

A 2 (group: PLC/PLC2)*2 (condition: social/non-social)*2 (task: pro-/anti-saccade) mixed ANOVA on latency was conducted on error rate to examine the social effects on attention processing between two PLC groups. Results showed a significant main effect of condition ($F_{1, 70} = 25.00$, $p < 0.001$, $\eta_p^2 = 0.26$) and a main effect of task ($F_{1, 70} = 125.18$, $p < 0.001$, $\eta_p^2 = 0.64$) with higher error rates for social than non-social stimuli (social: Mean ± SEM = 7.38% ± 0.54%, non-social: Mean ± SEM = 4.91% ± 0.51%) and increased error rates during anti-saccade than pro-saccade condition (anti: Mean ± SEM = 10.05% ± 0.76%, pro: Mean ± SEM = 2.24% ± 0.31%). The interaction between condition and task was also significant ($F_{1, 70} = 16.13$, $p <$

0.001, $\eta_p^2 = 0.19$), Post-hoc Bonferroni-corrected tests showed higher error rates for social compared to non-social stimuli during the anti-saccade condition (social: Mean ± SEM = 12.26% ± 0.88%, non-social: Mean ± SEM = 7.85% ± 0.90%, $p < 0.001$) but not the pro-saccade condition (social: Mean ± SEM = 2.51% ± 0.38%, non-social: Mean ± SEM = 1.96% ± 0.32%, $p = 0.11$).

To further explore whether there were significant differences between two PLC groups in the emotion-specific effects on error rate, a 2 (group: PLC/PLC2)*2 (task: pro-/anti-saccade)*6 (stimuli: angry/sad/fearful/happy/neutral/shape) mixed ANOVA was conducted. Results showed a main effect of task ($F_{1, 70} = 157.19$, $p < 0.001$, $\eta_p^2 = 0.69$) and a main effect of stimuli ($F_{5, 350} = 7.91$, $p < 0.001$, $\eta_p^2 = 0.10$). Examination of the main effect of task revealed higher error rates during anti-saccade condition compared to pro-saccade condition (anti: Mean ± SEM = 11.52% ± 0.81%, pro: Mean ± SEM = 2.38% ± 0.33%). Examination of the main effect of stimuli revealed higher error rates for faces compared to shapes (all *ps* < 0.006). Additionally, we found a significant interaction between stimuli and task ($F_{5, 350} = 5.12$, $p < 0.001$, $\eta_p^2 = 0.07$). Post-hoc Bonferroni-corrected comparisons revealed increased error rates for faces compared to shapes (all *ps* < 0.026) during anti-saccade rather than pro-saccade condition (all *ps* > 0.90). Together these results suggested that there were no significant differences between the two placebo groups for the social- and emotion-specific effects on attentional processing.

**Potential confounders**

To control for the potential confounding effects of between-subject differences in mood and personality traits, subjects were required to complete the following validated psychometric questionnaires including Positive and Negative Affect Schedule (PANAS), State-Trait Anxiety

Inventory (STAI), Liebowitz Social Anxiety Scale (LSAS, Heimberg et al., 1999), Social Interaction Anxiety Scale (SIAS, Mattick and Clarke, 1998), Beck Depression Inventory II (BDI-II, Beck et al., 1996), Autism Spectrum Quotient (ASQ, Baron-Cohen et al., 2001), Childhood Trauma Questionnaires (CTQ, Bernstein et al., 1998), Behavioral Inhibition System and Behavioral Activation System Scale (BIS/BAS, Carver and White, 1994), Action Control Scale (ACS, Kuhl, 1994), Cognitive Emotion Regulation Questionnaires (CERQ, Garnefski et al., 2001), 20-item Toronto Alexithymia Scale (TAS-20, Bagby et al., 1994).

**Table S1**. Demographics and questionnaire scores for subjects in PLC and AVP group included in the final analysis

| | PLC | AVP | t-value | p-value |
|---|---|---|---|---|
| Gender | 39 males | 39 males | | |
| Age (Mean ± SEM) | 22.74 ± 0.38 | 21.23 ± 0.30 | | |
| **Pre-task** | | | | |
| Positive and Negative Affect Schedule (PANAS) | | | | |
| Positive | 27.36 ± 0.98 | 28.21 ± 0.97 | 0.61 | 0.54 |
| Negative | 15.67 ± 0.79 | 15.67 ± 0.98 | 0.00 | 0.99 |
| State-Trait Anxiety Inventory (STAI) | | | | |
| SAI | 39.31 ± 1.54 | 40.05 ±1.43 | 0.35 | 0.72 |
| TAI | 40.23 ±1.31 | 39.82 ± 1.21 | 0.23 | 0.82 |
| Liebowitz Social Anxiety Scale (LSAS) | | | | |
| Avoid | 19.36 ± 1.66 | 18.36 ± 1.54 | 0.44 | 0.66 |
| Fear | 22.77 ± 2.01 | 22.67 ± 1.74 | 0.04 | 0.97 |
| Beck Depression Inventory (BDI-II) | 8.59 ± 1.09 | 7.59 ± 1.01 | 0.67 | 0.53 |
| Social Interaction Anxiety Scale (SIAS) | 52.10 ± 2.14 | 51.36 ± 1.71 | 0.27 | 0.79 |
| Autism Spectrum Quotient (ASQ) | 21.77 ± 0.98 | 20.38 ± 0.89 | 1.04 | 0.30 |
| Childhood Trauma Questionnaires (CTQ) | 40.79 ± 1.14 | 40.46 ± 1.34 | 0.19 | 0.85 |
| 20-item Toronto Alexithymia Scale (TAS-20) | 48.03 ± 1.61 | 50.31 ± 1.18 | 1.14 | 0.26 |
| Behavioral Inhibition System and | | | | |

| | | | | |
|---|---|---|---|---|
| Behavioral Activation System Scale (BIS/BAS) | | | | |
| BAS – Reward Responsiveness | 6.64 ± 0.25 | 6.97 ± 0.27 | 0.92 | 0.36 |
| BAS - Drive | 8.15 ± 0.37 | 7.77 ± 0.31 | 0.79 | 0.43 |
| BAS – Fun Seeking | 10.15 ± 0.39 | 10.33 ± 0.39 | 0.33 | 0.74 |
| BIS – Behavioral Inhibition | 16.21 ± 0.42 | 15.56 ± 0.46 | 1.03 | 0.31 |
| Action Control Scale (ACS) | | | | |
| Failure | 5.77 ± 0.51 | 5.28 ± 0.50 | 0.68 | 0.50 |
| Decision | 6.69 ± 0.54 | 6.69 ± 0.42 | 0.00 | 0.99 |
| Performance | 8.33 ± 0.31 | 8.77 ± 0.32 | 0.97 | 0.33 |
| Cognitive Emotion Regulation Questionnaires (CERQ) | 48.10 ± 1.19 | 48.38 ± 1.04 | 0.18 | 0.86 |
| **Post-task** | | | | |
| Positive and Negative Affect Schedule (PANAS) | | | | |
| Positive | 25.95 ± 1.19 | 28.15 ± 1.07 | 1.38 | 0.17 |
| Negative | 13.54 ± 0.77 | 13.95 ± 0.74 | 0.38 | 0.70 |
| State Anxiety Inventory (SAI) | 38.28 ± 1.47 | 37.62 ±1.09 | 0.36 | 0.72 |

TRANSPARENT REPORTING of TRIALS

CONSORT 2010 Flow Diagram

**Intranasal vasopressin like oxytocin increases social attention by influencing top-down control, but additionally enhances bottom-up control**

Qian Zhuang[1,#], Xiaoxiao Zheng[1,#], Benjamin Becker[1], Wei Lei[2], Xiaolei Xu[1,*], Keith M. Kendrick[1,*]

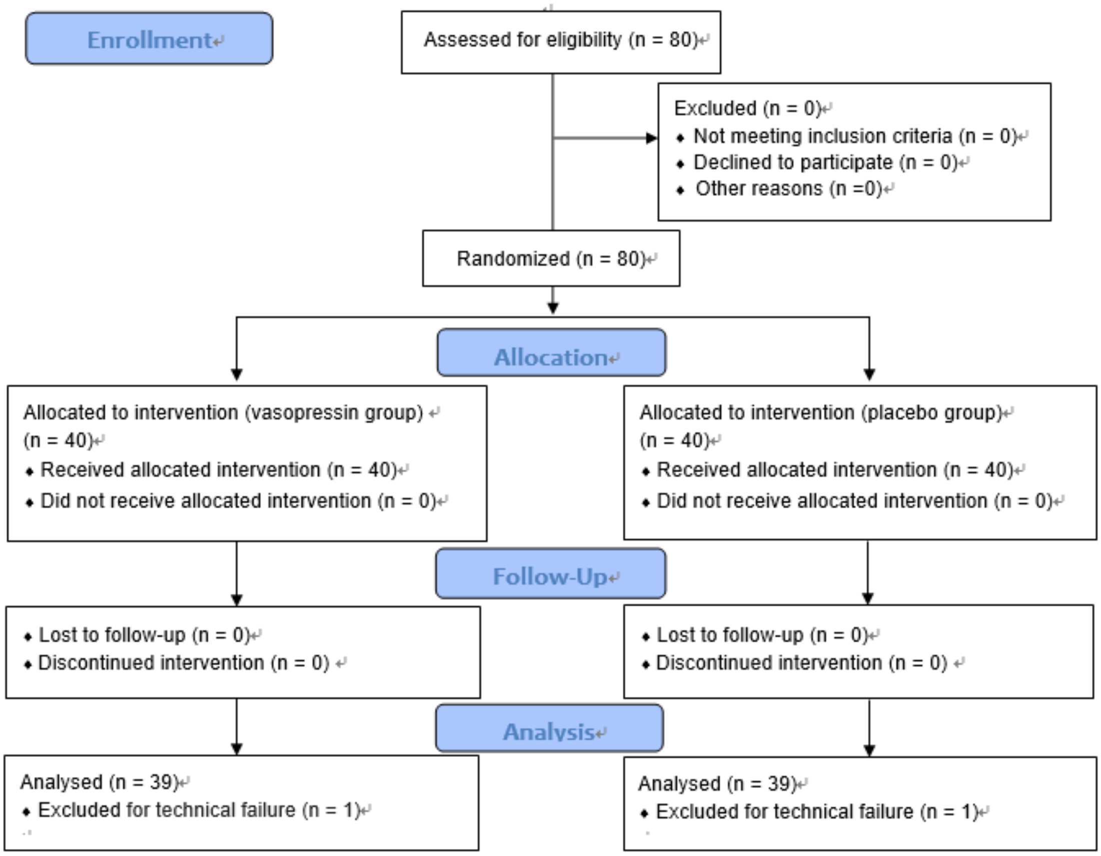


**Figure S1.** Consolidated standards of reporting trials (CONSORT) flow chart.
